# Distinct waking states for strong evoked responses in primary visual cortex and optimal visual detection performance

**DOI:** 10.1101/437681

**Authors:** Garrett T. Neske, David A. McCormick

## Abstract

Variability in cortical neuronal responses to sensory stimuli and in perceptual decision making performance is substantial. Moment-to-moment fluctuations in waking state or arousal can account for much of this variability. Yet, the nature of this variability across the full spectrum of waking states is often not completely characterized, leaving the characteristics of the optimal state for sensory processing unresolved. Using pupillometry in concert with extracellular multiunit and intracellular whole-cell recordings, we found that the magnitude and reliability of visually evoked responses in primary visual cortex (V1) of awake, passively behaving male mice increase as a function of arousal and are largest during sustained locomotion periods. During these high-arousal, sustained locomotion periods, cortical neuronal membrane potential was at its most depolarized and least variable. Contrastingly, behavioral performance of mice on two distinct visual detection tasks was generally best at a range of intermediate arousal levels, but worst during locomotion. These results suggest that large, reliable responses to visual stimuli in V1 occur at a distinct arousal level from that associated with optimal visual detection performance. Our results clarify the relation between neuronal responsiveness and the continuum of waking states, and suggest new complexities in the relation between primary sensory cortical activity and behavior.

## Introduction

Sensory-evoked responses of cortical neurons are known to exhibit substantial variability (Tolhurst et al., 1983; Shadlen and Newsome, 1998), a portion of which can be accounted for by baseline cortical network state (Arieli et al., 1996; Fontanini and Katz, 2008; Goris et al., 2014). While the most prominent cortical state change occurs upon the transition from sleep to wakefulness (Steriade, 2000; Steriade et al., 2001), the waking period itself is characterized by frequent, and often underappreciated state changes (McGinley et al., 2015b). These waking state fluctuations can profoundly impact sensory-evoked cortical responses and sensory-guided behaviors (Beaman et al., 2017; Engel et al., 2016; Pinto et al., 2013; Sachidhanandam et al., 2013; Speed et al., 2018). In the awake, head-fixed mouse preparation, a frequently used experimental paradigm of waking state fluctuation has been the comparison between periods of stillness and locomotion, particularly in primary visual cortex (V1). A broad consensus is that, compared to stillness or quiet wakefulness, locomotion enhances the visually evoked responses of V1 neurons while generally leaving neuronal tuning properties unaffected, a hallmark of a change in neuronal response gain (Bennett et al., 2013; Dadarlat and Stryker, 2017; Fu et al., 2014; Lee et al., 2014; Mineault et al., 2016; Niell and Stryker, 2010; Pakan et al., 2016; Polack et al., 2013).

The widespread finding of enhancement of V1 sensory responses during locomotion raises a question about the state dependence of cortical sensory processing more generally: how do state fluctuations occurring during non-locomotive wakefulness modulate cortical sensory responsiveness? While the bipartite, still-vs.-locomotion distinction has served as a tractable approach to investigate the state dependence of sensory processing in mice, this method leaves a large portion of the waking state space potentially disregarded. Using pupil diameter as an index of cortical state, several recent investigations have begun to address waking state changes occurring in the absence of locomotion. In the mouse visual system, increases in arousal without locomotion (as indexed by pupil dilation) have been shown to enhance the sensory-evoked signal-noise ratio and population encoding capacity of V1 neurons while decreasing their spontaneous firing rates (Reimer et al., 2014; Vinck et al., 2015). In the mouse auditory system, McGinley et al. (2015a) analyzed spontaneous and sound-evoked A1 activity as a function of a continuum of waking states, finding that evoked responses were largest and most reliable during intermediate arousal without locomotion, a state corresponding to low spontaneous neuronal activity levels and hyperpolarized membrane potential (V_m_) in A1 cortical neurons. The characteristics of spontaneous and evoked activity in mouse V1 as a continuous function of arousal have not been well explored.

Despite the significant interest in uncovering the characteristics and mechanisms of state-dependent sensory processing in mice, it is striking that very few studies have investigated the consequences of waking state modulations on sensory-guided behaviors. Only two previous studies using the head-fixed, locomoting mouse preparation have specifically addressed how waking state influences perceptual task performance. Bennett et al. (2013) showed that, compared with stillness, locomotion slightly, but significantly improved visual detection task performance. McGinley et al. (2015a), analyzing waking state as a continuum, showed that auditory detection task performance was optimal during intermediate arousal without locomotion. The optimal arousal level on the waking state spectrum for visual detection performance in mice is unknown.

Here we have studied the properties of spontaneous and visually evoked neuronal activity in awake, passively behaving mouse V1 as a function of the full waking state spectrum by using pupil size as a continuous index of arousal. We found that, while spontaneous firing of V1 neurons is minimal at intermediate arousal without locomotion, visually evoked response magnitude and reliability are monotonic increasing functions of arousal, and highest during periods of sustained locomotion. Whole-cell patch clamp recordings revealed that this monotonic increase in sensory responsiveness is associated with a corresponding depolarization of V1 neuronal V_m_ and decrease in V_m_ variance. Contrastingly, we found that performance of mice on visual detection tasks was generally best during a range of intermediate arousal levels, but suboptimal during locomotion. Our results provide a more complete picture of the modulation of sensory cortical activity and sensory-guided behavior by waking state.

## Results

### Waking state, as indexed by pupil size, varies over a wide range

While the relation between pupillary dilation/constriction and arousal and other cognitive variables has been investigated for well over half a century (Goldwater, 1972; Loewenfeld, 1999), several recent studies in awake, head-fixed mice have detailed a very close relationship between pupil size and cortical network state, even at the level of cortical membrane potential (Reimer et al., 2014; Vinck et al., 2015; McGinley et al., 2015a,b).

Here we also used pupil size as a continuous gauge of state to study the properties of V1 neuronal responses and visually guided behavior across the full waking state spectrum. In our study, mice were head-fixed and free to walk on a cylindrical treadmill while an LCD monitor presented visual stimuli to the left visual hemifield and a camera captured video of the right eye, illuminated by an IR LED (Fig. 1a). As in McGinley et al. (2015a), we found that waking state, as indexed by pupil size, varied over a wide range during a typical recording session lasting 1-2 hrs (Fig. 1b). To quantify pupil size, we calculated the diameter of the best-fit circle to the pupil in individual video frames (pupil diameter) (see Materials and Methods, “Quantification of pupil size”). All pupil diameters reported here are normalized to the maximal pupil diameter recorded during a session, which always occurred during sustained locomotion bouts. Normalized pupil diameter was approximately normally distributed across animals and recording sessions (Fig. 1c). Since we used both passively behaving and task-engaged animals in our study, we constructed pupil diameter probability distributions for these two categories separately (Fig. 1c). Notably, pupil diameter distributions were centered on different values for passively behaving and task-engaged animals; while distributions for the latter were centered on pupil diameters ~70% constricted from the maximum, distributions for the former were centered on pupil diameters ~50% constricted from the maximum (Fig. 1c).

**Figure 1.**
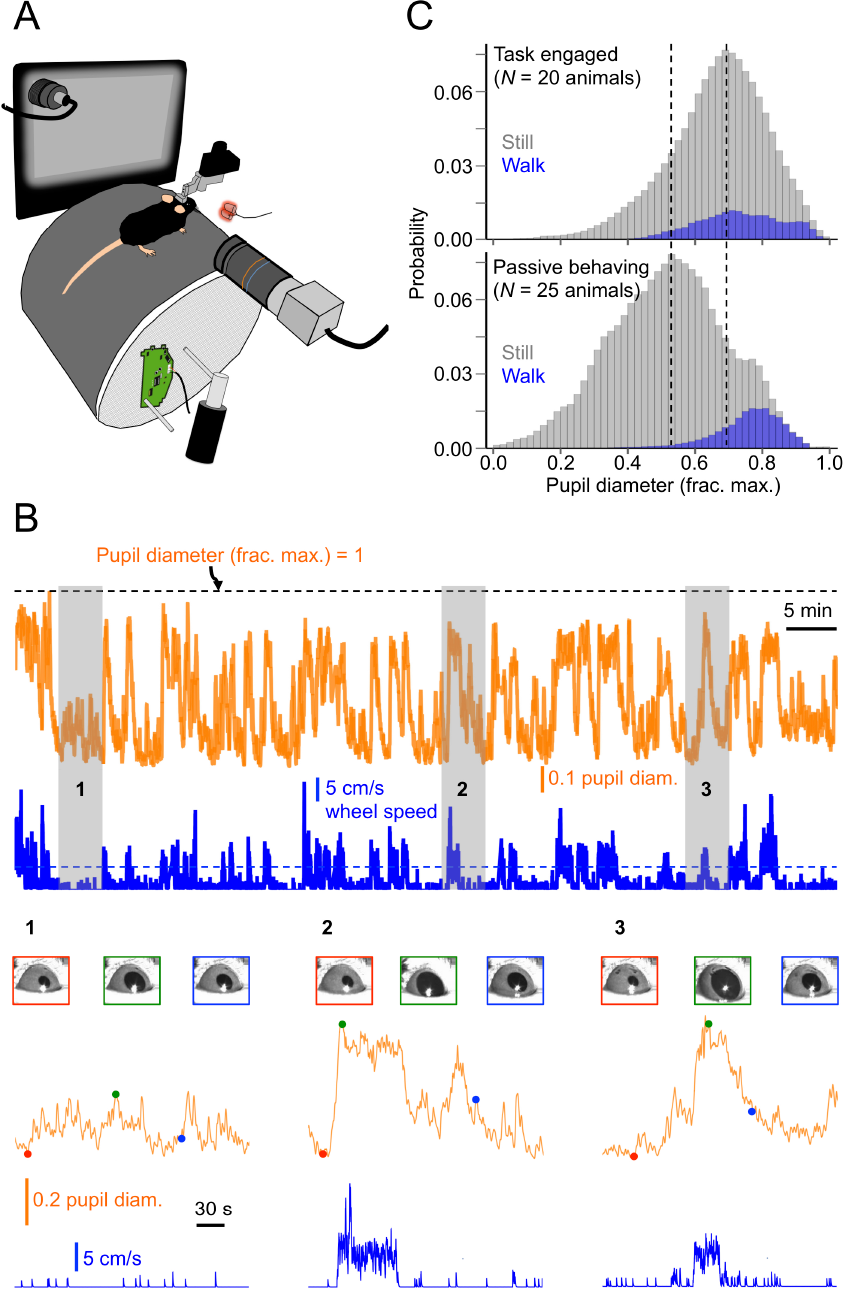
Waking state, as indexed by pupil size, varies over a wide range. **(A)** Basic experimental set-up for the head-fixed mouse preparation. An LCD monitor is mounted to the mouse’s left, presenting visual stimuli to the left eye, and a USB camera is mounted to the mouse’s right, capturing video of the right eye, which is illuminated by 2 IR LEDs. A photodiode mounted to the upper-left corner of the LCD monitor gives precise timestamps of stimulus onsets. An optical mouse mounted to the side of the wheel acquires wheel speed. **(B)** Representative traces of normalized pupil diameter and locomotion speed for an entire example recording session (~90 min). The horizontal dashed line plotted with wheel speed indicates the 5 cm/s threshold for qualifying as a locomotion bout (see Materials and Methods, “State-dependent electrophysiological metrics”). Insets magnify the numbered regions and show regions of interest (ROIs) around the eye for selected times. **(C)** Normalized pupil diameter distributions, sorted according to whether mice were still or walking, for all task-engaged (*N* = 20) or passively behaving (*N* = 25) animals in the data set. Vertical dotted lines indicate the peaks of the two distributions.

### Spontaneous V1 spiking activity is minimal at intermediate arousal without locomotion

We first considered how spontaneous spiking activity in V1 is modulated as a function of arousal in passively behaving animals. We recorded spontaneous multi-unit activity (MUA) and LFP primarily in layer 5 V1 (Fig. 2a) and binned this activity as a function of pupil diameter (see Materials and Methods, “State-dependent electrophysiological metrics”) (N = 17 animals, n = 23 recordings). Note that pupil diameter bin widths are chosen such that an equal amount of data falls into each bin, making the bin widths for the smallest and largest pupil diameters often larger than those for intermediate pupil diameters. We also dissociated the data into periods in which the mice were still and periods in which they were engaged in sustained locomotion bouts (see Materials and Methods, “State-dependent electrophysiological metrics”). Spontaneous activity was high at the lowest arousal levels (smallest pupil diameter) and decreased with increased arousal, reaching a minimum at still periods with intermediate arousal. After this intermediate-arousal state in which spontaneous activity was minimal, increased arousal resulted in increased spontaneous activity, increasing even further during locomotion (Fig. 2b). We also observed this U-shaped function of spontaneous activity versus arousal when separately analyzing MUA from layer 2/3 (Fig. 2c).

Note that in Fig. 2b,c, and other similar figures throughout this study, measured baseline pupil diameter accounts for the lagged temporal relationship between pupil diameter and electrophysiological variables (see Materials and Methods, “State-dependent electrophysiological metrics”). Not accounting for this lag, however, did not significantly change the essential relationships between baseline pupil diameter and V1 cortical activity (Fig. 2-1).

**Figure 2.**
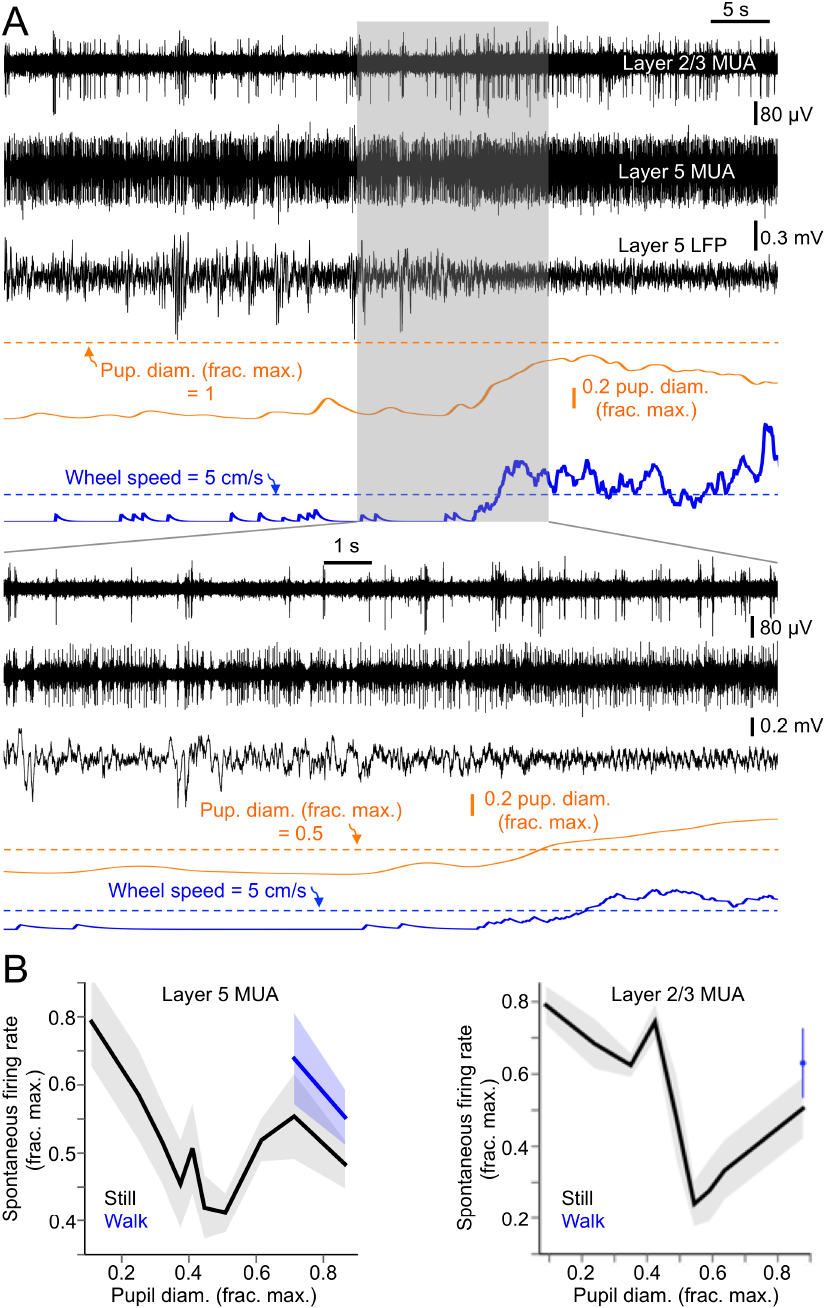
Spontaneous firing rates in V1 are minimal during intermediate arousal without locomotion. **(A)** Example traces of multi-unit activity (MUA) simultaneously recorded in layer 2/3 and 5 of V1, along with layer 5 LFP, normalized pupil diameter, and locomotion speed. **(B)** Spontaneous firing rates (as a fraction of the maximum spontaneous firing rate recorded during a session) in layer 5 and 2/3 as a function of baseline pupil diameter and sorted by locomotion status. Baseline pupil diameter bin widths were chosen such that an equal amount of data fell into each bin. (Layer 5: N = 17 animals, n = 23 recordings; Layer 2/3: N = 6 animals, n = 6 recordings. Data are mean ± 68% CI).

**Figure 2-1.**
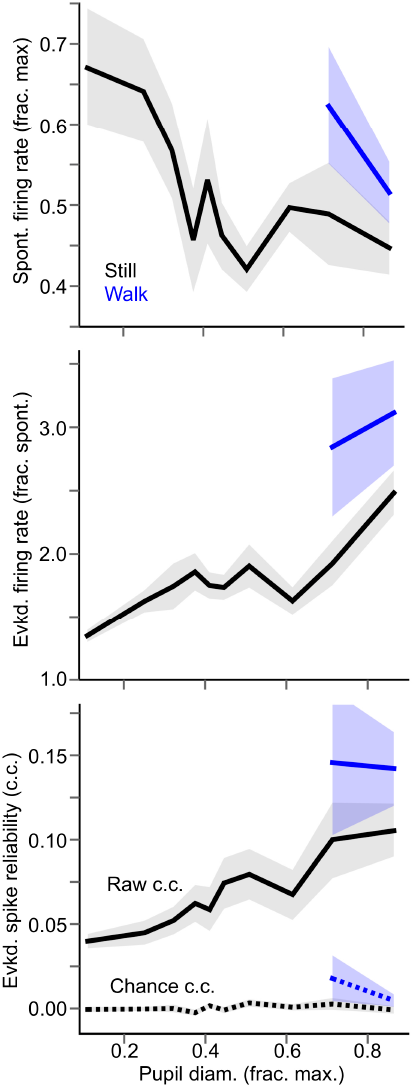
State-dependent spontaneous and evoked layer 5 V1 activity, not accounting for lag between pupil diameter and cortical activity. Top plot as in Fig. 2b, and bottom 2 plots as in Fig. 3b-c. Here, however, baseline pupil diameter was calculated as the average normalized pupil diameter 500 ms before the spontaneous or evoked epoch analyzed without taking into account the lagged relationship between changes in pupil diameter and changes in V1 spiking activity. Not accounting for this lag did not affect the finding that spontaneous activity is minimal at intermediate arousal without locomotion and that evoked responses are enhanced monotonically with arousal. For pupil diameter bins, bin widths were chosen such that an equal amount of data fell into each bin.

### Sensory evoked V1 spiking responses are monotonically enhanced with arousal

Visually evoked responses of neurons in mouse V1 are known to be enhanced by locomotion (Bennett et al., 2013; Dadarlat and Stryker, 2017; Fu et al., 2014; Lee et al., 2014; Mineault et al., 2016; Niell and Stryker, 2010; Pakan et al., 2016; Polack et al., 2013). Furthermore, increases in arousal without locomotion have also been shown to enhance V1 neuronal responsiveness and sensory encoding capacity (Reimer et al., 2014; Vinck et al., 2015). How do visually evoked responses in mouse V1 vary across the continuum of arousal, and which level of arousal within this continuum is associated with the largest and most reliable visual responsiveness? To address this question we quantified visually evoked MUA responses in V1 layer 5 as a function of baseline pupil diameter in passively behaving animals (N = 17 animals, n = 23 recordings). To maximize the size of the neuronal population contributing to the recorded MUA, we used full-contrast Gaussian noise movies as visual stimuli (Fig. 3a; see Materials and Methods, “Visual stimulus presentation”), which approximately match the spatial frequency spectrum of the stimulus to the distribution of spatial frequency preferences of V1 neurons (Niell and Stryker, 2008).

We found that the magnitude of evoked V1 multi-unit spiking responses (as a fraction of baseline spiking activity, i.e., signal-noise ratio) and the reliability of these responses were monotonic increasing functions of arousal, with the largest and most reliable responses occurring during locomotion (Fig. 3b,c; Fig. 2-1). Notably, locomotion itself enhanced response magnitude and reliability beyond that associated with high arousal during still periods; when compared within bins of large pupil diameter (i.e. pupil diameter bins in which locomotion occurred), locomotion significantly increased signal-noise ratio compared with stillness (Fig. 3d) (signal-noise ratio at large pupil diameters during stillness vs. signal-noise ratio at large pupil diameters during walking: 1.98 ± 0.16 vs. 2.83 ± 0.36, p = 0.02, rank-sum test). Response reliability was also significantly enhanced during locomotion compared with stillness (Fig. 3e) (reliability at large pupil diameters during stillness vs. reliability at large pupil diameters during walking: 0.076 ± 0.02 vs. 0.12 ± 0.02, p = 0.00009, rank-sum test). Relatedly, in individual recordings, we determined that the largest (Fig. 3f) and most reliable (Fig 3g) evoked responses occurred most frequently in the bin associated with the largest pupil diameters, both during stillness (Fig. 3f,g, top) and especially when locomotion bouts were included in the analysis (Fig. 3f,g, bottom). We also observed monotonic-increasing V1 sensory response enhancements as a function of arousal when analyzing layer 2/3 MUA separately (Fig. 3-1).

**Figure 3.**
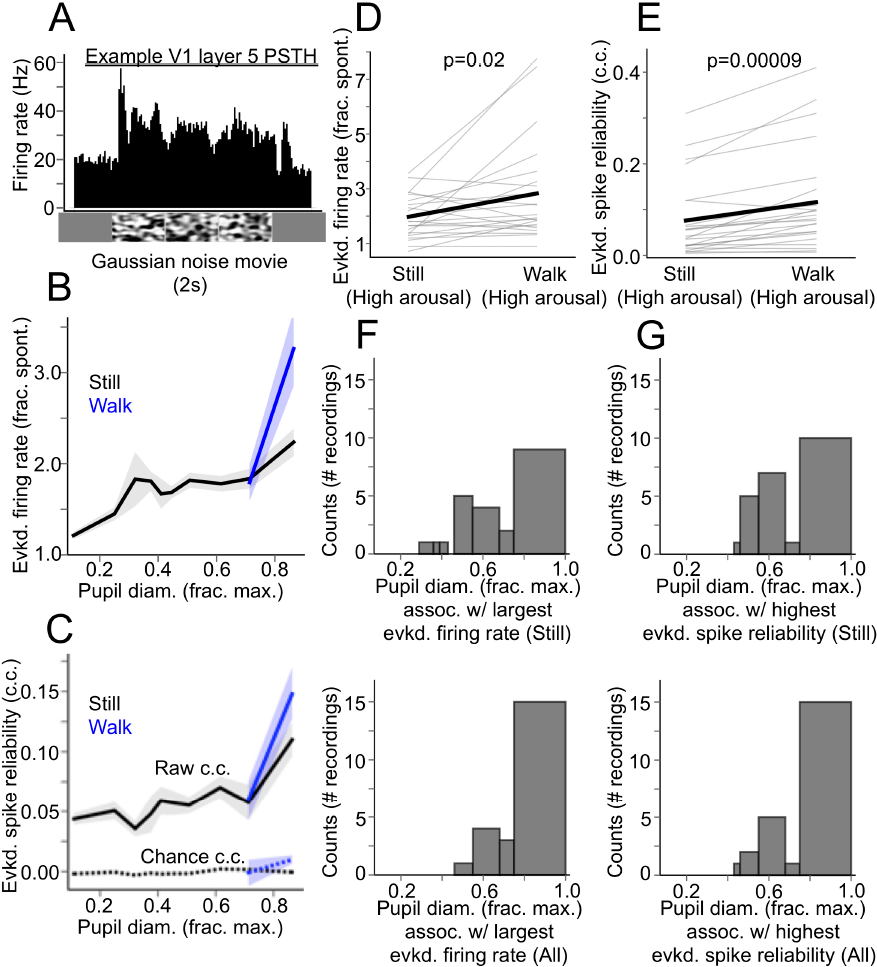
Visually evoked spiking responses in V1 are enhanced monotonically with increasing arousal, and largest during locomotion. **(A)** Example peri-stimulus time histogram (PSTH) of the evoked layer 5 MUA response to a full-contrast Gaussian noise movie. **(B)** Evoked layer 5 multi-unit firing rate (as a fraction of spontaneous, baseline firing rate 500 ms before stimulus presentation) in response to full-contrast Gaussian noise movies, as a function of baseline pupil diameter, and sorted by locomotion status. **(C)** Trial-by-trial reliability (cross-correlation, c.c.) of multi-unit spiking responses to full-contrast Gaussian noise movies as a function of baseline pupil diameter, and sorted by locomotion status. “Raw” denotes the pairwise cross-correlation between evoked spiking responses to the same Gaussian noise movie, and “chance” denotes the pairwise crosscorrelation between evoked spiking responses and periods of spontaneous spiking activity occurring in the same pupil diameter bin, a correction for cross-correlation increases due to increased spiking alone. **(D)** Within-recording comparisons of evoked layer 5 multi-unit firing rate (fraction of baseline firing rate) between still and locomotion periods during high arousal (i.e. in pupil diameter bins in which locomotion occurred). P-value from rank-sum test. **(E)** Within-recording comparisons of evoked layer 5 multi-unit spike reliability (trial-by-trial crosscorrelations of evoked PSTHs) between still and locomotion periods during high arousal. P-value from rank-sum test. **(F)** (*Top*) Histogram of extracellular MUA recordings, sorted into bins of pupil diameter (during stillness) in which the largest evoked responses occurred during an individual recording. (*Bottom*) Same histogram, but including locomotion periods. (*G*) As in F, but for highest evoked spike reliability. For pupil diameter bins in *B, C, F, and G*, bin widths were chosen such that an equal amount of data fell into each bin. (N = 17 animals, *n* = 23 recordings. Data are mean ± 68% CI).

**Figure 3-1.**
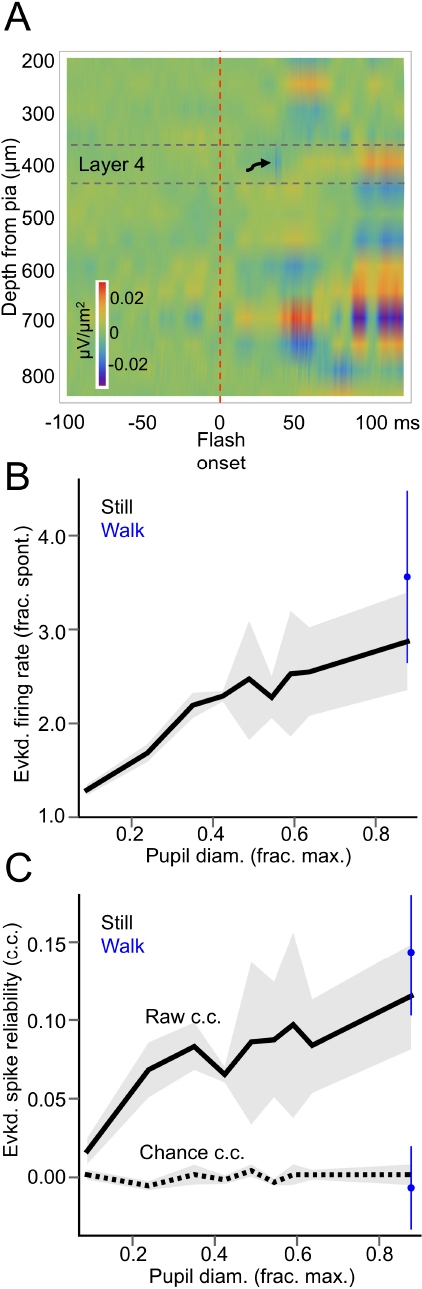
State-dependence of evoked responses in V1 layer 2/3 is similar to that of layer 5. **(A)** Representative current-source density (CSD) plot used to localize silicon probe contacts residing in layer 2/3. Within ~40 ms of a 50-ms full-screen flash, a strong, short-latency sink is evident in mid-layers (arrow), followed by delayed sinks in more superficial and deep layers after 50 ms. Even stronger sinks are evident in putative deep layer 5 100 ms after stimulus onset, likely due to polysynaptic activity induced by the stimulus. Contacts in layer 2/3 were considered to be 50 - 100 μm above the estimated layer 4 boundary. **(B)** Evoked layer 2/3 multiunit firing rate (as a fraction of spontaneous, baseline firing rate 500 ms before stimulus presentation) in response to full-contrast Gaussian noise movies, as a function of baseline pupil diameter, and sorted by locomotion status. **(C)** Trial-by-trial reliability (cross-correlation, c.c.) of layer 2/3 multi-unit spiking responses to full-contrast Gaussian noise movies as a function of baseline pupil diameter, and sorted by locomotion status. “Raw” denotes the pairwise cross-correlation between evoked spiking responses to the same Gaussian noise movie, and “chance” denotes the pairwise cross-correlation between evoked spiking responses and periods of spontaneous spiking activity occurring in the same pupil diameter bin, to correct for cross-correlation increases due to increased spiking alone. For pupil diameter bins in *B* and C, bin widths were chosen such that an equal amount of data fell into each bin. (*N* = 6 animals, *n* = 6 recordings. Data are mean ± 68% CI).

The monotonic increase in V1 sensory responsiveness with arousal that we have described could be due in part to the simultaneous monotonic increase in pupil size itself, perhaps by allowing more light to enter the eye. To ensure that the pupil-size-dependent effects on V1 neuronal responses were due to state changes and not simply due to changes in pupil size, we recorded visually evoked MUA responses in a separate cohort of animals (N = 17 animals, n = 17 recordings) in which the pupil of the left, visually stimulated eye was kept maximally dilated throughout the entire recording session via application of atropine, while the pupil size of the right eye was free to continuously vary with state (see Materials and Methods, “Atropine experiments”). Even when the pupil of the visually stimulated eye was kept maximally dilated, V1 neuronal responses were still enhanced as a function of the pupil size of the non-visually stimulated eye (Fig. 3-2).

**Figure 3-2.**
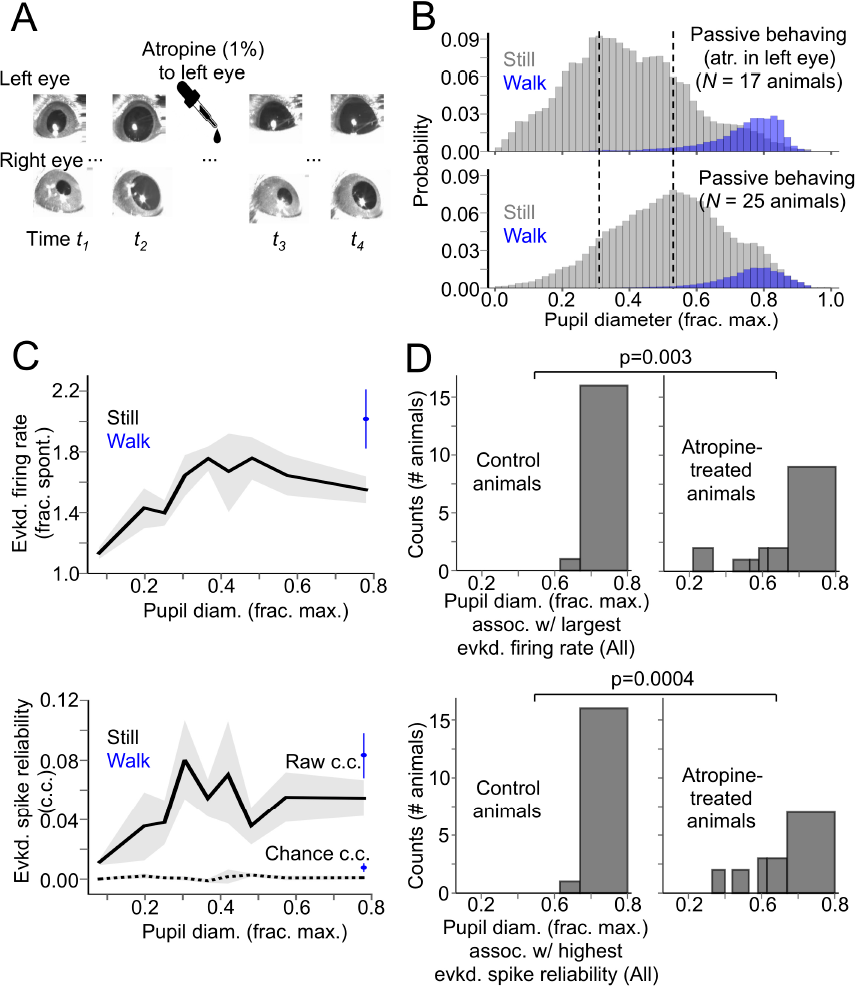
Complete artificial dilation of the visually stimulated eye does not fundamentally alter the enhancement of evoked V1 responses as a function of pupil size. **(A)** Individual video frames from sessions in which 2 cameras were positioned to image each eye. Prior to atropine application, the pupil sizes of both eyes fluctuate coherently. After atropine application to the left eye, the pupil of the right eye still varies with state, but the pupil of the left eye is always fully dilated. **(B)** Probability histogram of normalized pupil diameter for animals treated with atropine in the visually stimulated eye (*top*), compared with control animals under the same passive-behaving conditions (*bottom*, distribution same as in Fig. 1c). While normalized pupil diameter spans the full range in atropine-treated animals, the distribution is notably centered on smaller values. Vertical dotted lines indicate the peaks of the two distributions. **(C)** Evoked firing rate and spike reliability as a function of baseline pupil diameter and sorted by locomotion status for atropine-treated animals. **(D)** Histograms (counts = number of animals) of pupil diameter bins (still and locomotion periods combined) associated with the largest evoked firing rate recorded per animal (*top*) and the highest evoked spike reliability recorded per animal (*bottom*), compared between atropine-treated animals and control animals (control animals same as those reported in Fig. 3). P-values are from Fisher’s exact test. For pupil diameter bins in *C* and *D*, bin widths were chosen such that an equal amount of data fell into each bin. (*N* = 17 animals, *n* = 17 recordings. Data are mean ± 68% CI).

Notably, however, the distributions of pupil diameter bins associated with the largest and most reliable sensory-evoked responses in the atropine-treated animals were significantly different from those of control animals (Fig. 3-2d; response magnitude: p=0.003, response reliability: p=0.0004, Fisher’s exact test). Thus, while the majority of the pupil-size-dependent enhancement of sensory-evoked responsiveness described in Fig. 3 is most likely attributable to state changes, we do not rule out an additional, smaller contribution of pupil size per se.

**Figure 4.**
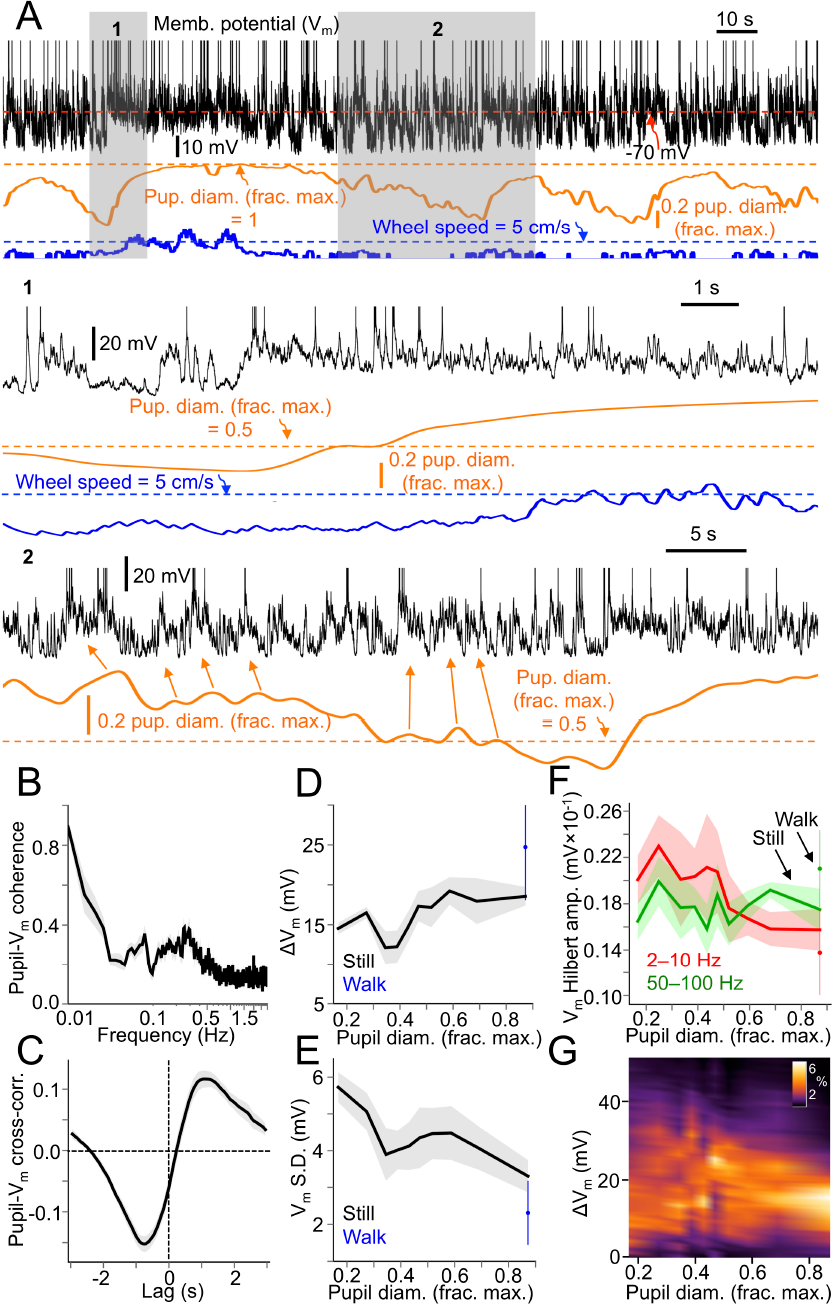
Membrane potential (V_m_) of V1 cortical neurons exhibits depolarization and decreased variability as a function of arousal, with most depolarized and least variable V_m_ during locomotion. **(A)** Example traces of V_m_ from a V1 layer 5 regular-spiking, putative pyramidal cell, pupil diameter, and locomotion speed. Inset 1 emphasizes V_m_ changes associated with the transition from stillness to locomotion. Inset 2 emphasizes the relationship between pupillary microdilations (arrows) and V_m_ depolarizations during stillness. Spikes are truncated in example traces to emphasize subthreshold behavior. **(B)** Coherence between pupil diameter and V_m_. Coherence is high at low frequencies (0.01-0.02 Hz), with a smaller secondary peak from ~0.1-0.3 Hz, likely reflecting the tracking of pupillary microdilations by V_m_. **(C)** Cross-correlation between pupil diameter and V_m_, showing a lag of ~1 s between changes in V_m_ and associated changes in pupil diameter. **(D)** Mean spontaneous V_m_ (expressed as a change in V_m_ from the most hyperpolarized V_m_ recorded during a session) as a function of baseline pupil diameter and sorted by locomotion status. **(E)** V_m_ standard deviation as a function of baseline pupil diameter and sorted by locomotion status. **(F)** V_m_ Hilbert amplitude at 2-10 Hz and 50-100 Hz as a function of baseline pupil diameter and sorted by locomotion status. **(G)** Density plot, pooling V_m_ values recorded from all cells in the data set, encapsulating data shown in *D* and E. Each vertical slice of the plot is the probability distribution of ΔV_m_ (the difference between V_m_ in a recorded cell and the most hyperpolarized V_m_ recorded in that cell) in a given pupil diameter range. A more dispersed probability distribution is evident at smaller pupil diameters, while a narrower distribution is evident at larger pupil diameters. For pupil diameter bins in *D - G*, bin widths were chosen such that an equal amount of data fell into each bin. (N = 8 animals, *n* = 10 cells. Data are mean ± 68% CI.).

### Membrane potential of V1 neurons becomes more depolarized and less variable with increased arousal

Transitions from stillness to locomotion have previously been shown to effect dramatic changes in the V_m_ of cortical neurons in mouse V1 (Bennett et al., 2013; Polack et al., 2013) and A1 (Schneider et al., 2014; Zhou et al., 2014); compared with stillness, locomotion is associated with a more depolarized V_m_ and lower V_m_ variance. The nature of cortical V_m_ dynamics across the continuum of arousal, however, has not been well studied. Using pupil size as a continuous index of arousal, McGinley et al. (2015a) showed that V_m_ in A1 neurons was most hyperpolarized and least variable during intermediate arousal without locomotion. We were interested in determining if similar V_m_ dynamics as a continuous function of arousal were also manifested in V1.

We performed whole-cell patch clamp recordings from regular-spiking, putative pyramidal cells in V1 layer 5 (average depth from pia: 497 (¼, S.D.: 39 ¼m) of awake, passively behaving mice while monitoring pupil diameter and locomotion (Fig. 4a) (N = 8 animals, n = 10 cells). V_m_ of V1 neurons was coherent with slow (0.01-0.02 Hz) fluctuations in pupil diameter (Fig. 4b), with a secondary peak in coherence between ~0.1-0.3 Hz, likely reflecting the tracking of pupillary microdilations by V_m_ (see also inset 2 in Fig. 4a).

**Figure 5.**
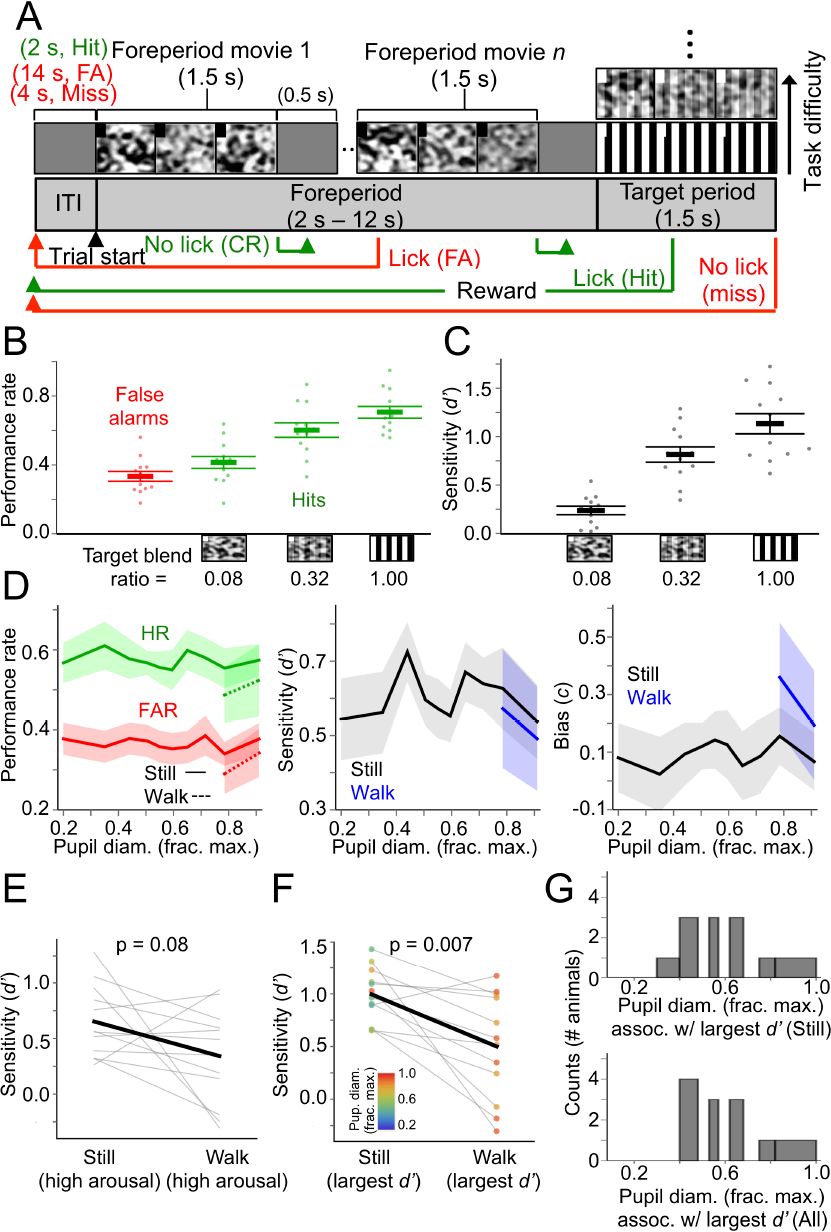
Performance of a target-in-noise visual detection task is suboptimal during locomotion. **(A)** Trial structure of the task. Each trial begins with a variable-duration foreperiod consisting of a sequence of Gaussian noise movies, during which the mouse must withhold licking. Mice must lick during the target period, in which a drifting square-wave grating is embedded in one of the noise movies with different blend ratios, modulating task difficulty. Note that for display purposes, the aspect ratio of the visual stimulus screen in the figure is different from that of the actual LCD monitor. **(B)** Overall performance rates (false alarm rates and hit rates) across animals for the different target levels presented during behavior sessions. **(C)** Overall perceptual sensitivities (d′) across animals for the different target levels presented during behavior sessions. **(D)** Performance rate, perceptual sensitivity (d′), and decision bias (c) as a function of baseline pupil diameter, and sorted by locomotion status. **(E)** Within-animal comparisons of d′ for stillness versus locomotion during high arousal (i.e. pupil diameter bins in which locomotion occurred). P-value is from rank-sum test. **(F)** Within-animal comparisons of the largest d’ prime recorded during stillness versus locomotion. Data points are colored according to the pupil diameter bin in which they were recorded. P-value is from rank-sum test. **(G)** Histograms (counts = number of animals) of the pupil diameter bin in which the largest d’ was recorded for each animal during stillness (top) and during stillness and locomotion combined (bottom). For pupil diameter bins in D and G, bin widths were chosen such that an equal amount of data fell into each bin. (N = 12 animals, n = 81 sessions. Data are mean ± 68% CI.).

Pupillary fluctuations lagged ~1 s behind associated fluctuations in Vm (Fig. 4c). Considering the relationship between baseline arousal (i.e. pupil diameter) and Vm characteristics, baseline Vm exhibited minimal change except during high arousal periods with locomotion (Fig 4d). Vm variability (measured as the standard deviation of baseline Vm) exhibited a progressive decrease as a function of increasing arousal, with the least variable Vm occurring during locomotion (Fig. 4e). Thus, a possible Vm correlate of the enhanced sensory responsiveness of V1 neurons as a function of arousal is a continual reduction of Vm variance. Furthermore, during the highest arousal levels, coinciding with sustained locomotion bouts, V1 neurons were at their most depolarized and least variable. Thus, the Vm correlate of the augmented sensory responsiveness of V1 cortical neurons during high arousal with locomotion is depolarization and reduced Vm variability.

### Visual detection performance is suboptimal during locomotion, and often optimal at intermediate arousal

The relationship between arousal and performance on demanding perceptual decision making tasks often follows a principle first articulated by Yerkes and Dodson (1908): task performance is optimal during intermediate arousal levels and degrades with either hypo- or hyperarousal. McGinley et al. (2015a) showed that performance of mice on an auditory detection task largely followed the Yerkes-Dodson inverted-U function of arousal. The arousal-dependence of auditory task performance was congruous with that of sound-evoked A1 responses in that the magnitude, reliability, and gain of these responses were also inverted-U functions of arousal, suggesting that the inverted-U relationship between arousal and performance might be due in part to activity in A1. Here we have shown, contrastingly, that the relationship between arousal and evoked responses in V1 is one of monotonic enhancement. What are the consequences of this relationship for performance of visual tasks in mice? To answer this question, we trained mice on two distinct visual detection tasks and quantified their performance as a function of pupil size and locomotion.

**Figure 6.**
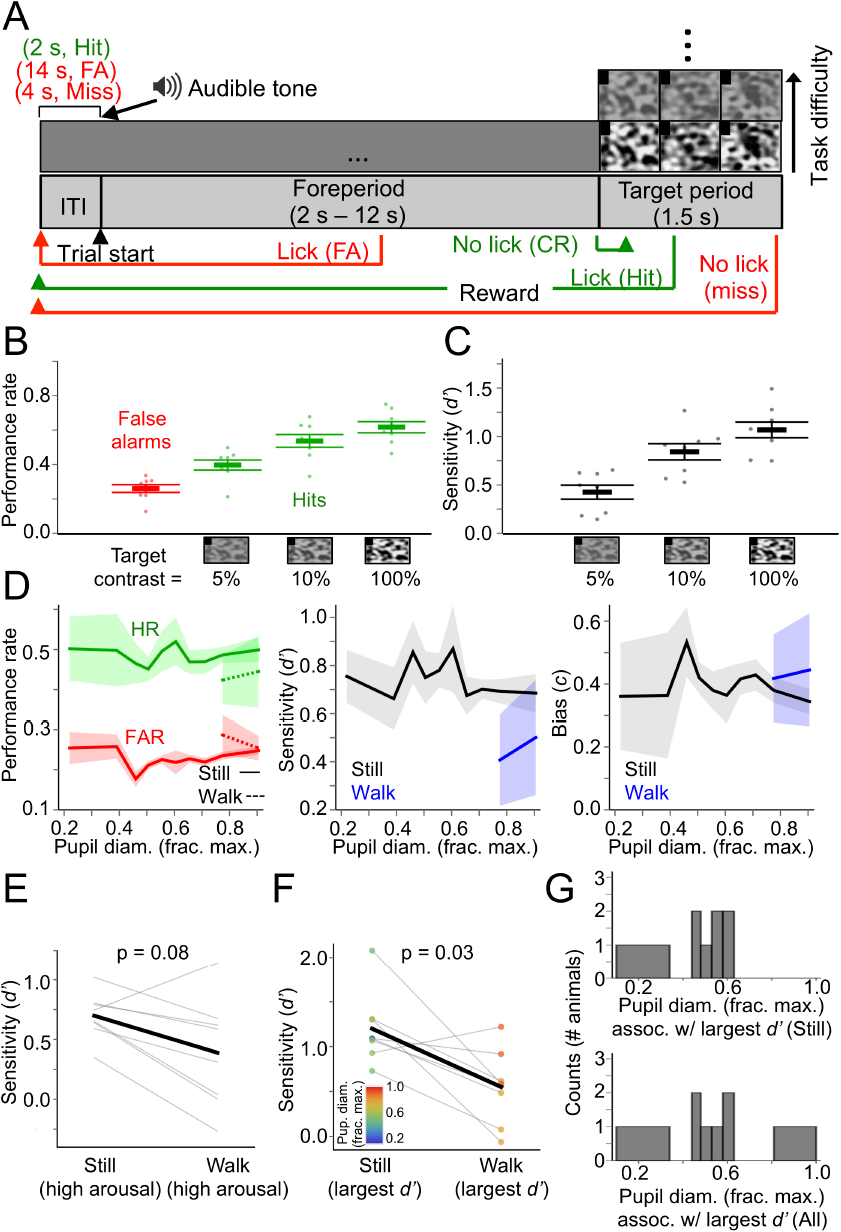
Visual detection of Gaussian noise movies is suboptimal during locomotion. **(A)** Trial structure of the task. Each trial begins with a variable-duration foreperiod (signaled by an audible tone) consisting of an isoluminant gray screen, during which the mouse must withhold licking. Mice must lick during the target period, in which a Gaussian noise movie is presented at different contrasts. Note that for display purposes, the aspect ratio of the visual stimulus screen in the figure is different from that of the actual LCD monitor. **(B)** Overall performance rates (false alarm rates and hit rates) across animals for the different target levels presented during behavior sessions. **(C)** Overall perceptual sensitivities (*d*′) across animals for the different target levels presented during behavior sessions. **(D)** Performance rate, perceptual sensitivity (*d*′), and decision bias (*c*) as a function of baseline pupil diameter, and sorted by locomotion status. **(E)** Within-animal comparisons of *d’* for stillness versus locomotion during high arousal (i.e. pupil diameter bins in which locomotion occurred). P-value is from rank-sum test. **(F)** Within-animal comparisons of the largest *d’* prime recorded during stillness versus locomotion. Data points are colored according to the pupil diameter bin in which they were recorded. P-value is from rank-sum test. **(G)** Histograms (counts = number of animals) of the pupil diameter bin in which the largest *d’* was recorded for each animal during stillness (*top*) and during stillness and locomotion combined (*bottom).* For pupil diameter bins in *D* and G, bin widths were chosen such that an equal amount of data fell into each bin. (N = 8 animals, *n* = 59 sessions. Data are mean ± 68% CI.).

In one task, mice (N = 12 animals, n = 81 sessions) were trained to lick a spout for water reward during a target period in which a drifting square-wave grating was blended with a Gaussian noise movie (hit), and withhold licking during a preceding, variable-length foreperiod that contained sequences of Gaussian noise movies without a blended grating (correct rejection) (Fig. 5a; Movie 1). This task is analogous in trial- and stimulus-structure to the auditory detection task used in McGinley et al. (2015a). Incorrect responses on this visual detection task include either licking during one of the foreperiod Gaussian noise movies (false alarm) or failing to lick during the grating-in-noise target period (miss). This target-in-noise detection task was psychometric, with gratings blended with noise movies at several different levels during individual sessions (see Materials and Methods, “Habituation to head fixation and behavioral training”; Fig. 5b,c; Fig. 5-1a). We found that locomotion was predominately a suboptimal state for task performance, whereas, for most animals, the optimal state for performance was one of intermediate arousal during stillness (Fig. 5d-g). On an animal-by-animal basis, comparing perceptual sensitivity (d′) to the target stimulus between still and locomotion periods during high arousal (i.e. pupil diameter bins in which locomotion occurred), we found that, with the exception of 3 out of 12 animals, locomotion was associated with smaller, though not significantly smaller d’ (Fig. 5e) (d’ for high arousal during stillness vs. d’ for high arousal during locomotion: 0.65 ± 0.03 vs. 0.34 ± 0.13, p = 0.08, rank-sum test). Comparing optimal (i.e. largest) d’ between still and locomotion periods showed that, with the exception of 2 out of 12 animals, d’ was significantly higher during still periods (Fig 5f) (d’ for still vs. locomotion: 1.00 ± 0.20 vs. 0.50 ± 0. 07, p = 0.007, rank-sum test). That is, for each animal, the largest d’ during still periods was almost always larger than the largest d’ during locomotion periods. While the arousal level (i.e. pupil diameter bin) corresponding to the largest d’ was variable across animals, this level most frequently corresponded with pupil diameters ~40-60 % constricted from the most dilated pupil diameter (Fig. 5g). Considering still periods alone, 10 out of 12 animals performed the task with the largest d’ when pupil diameter was ~30-70% constricted from the maximum, while only 2 out of 12 animals performed the task optimally during high arousal periods (~80-100% pupil diameter) (Fig. 5g, top). This distribution of optimal performance vs. pupil diameter was similar when locomotion periods were included in the analysis (Fig. 5g, bottom). Taken together, these results suggest that performance on a target-in-noise visual detection task is suboptimal during high arousal with locomotion and that a wide range of intermediate-arousal, non-locomotive states are compatible with task performance.

**Figure 5-1.**
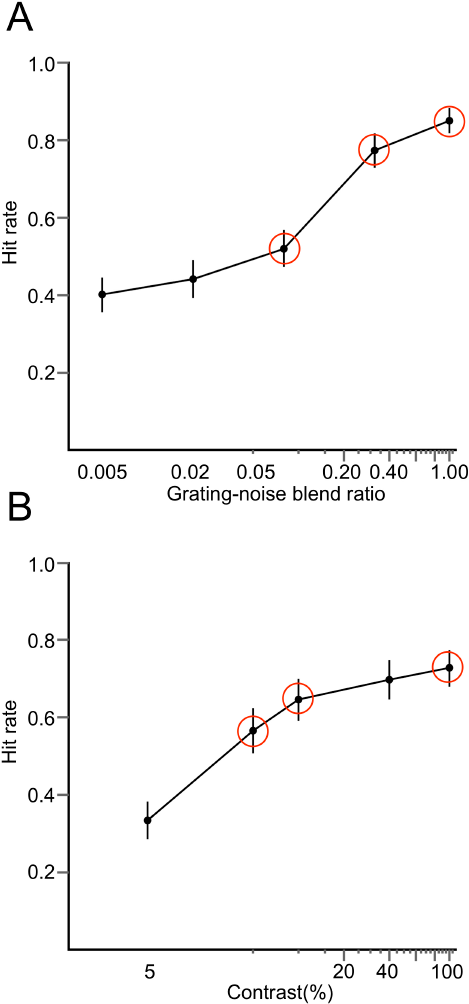
Psychometric data for visual detection tasks. **(A)** Psychometric curve for the target-in-noise detection task, averaged over all animals and sessions for which target stimuli were presented with 5 different grating-noise blend ratios (N = 12 animals, *n* = 170 sessions, separate from the sessions in which pupillometry and locomotion data were collected). Circled blend ratios were the 3 target levels used during sessions in which pupillometry and locomotion data were collected. **(B)** As in A, but for the noise detection task (N = 8 animals, *n* = 116 sessions, separate from the sessions in which pupillometry and locomotion data were collected). (Data are mean ± 68% CI.).A

We designed our target-in-noise visual detection task primarily to facilitate comparisons of state-dependent task performance with that of a comparably designed auditory detection task (McGinley et al., 2015a). The target-in-noise visual detection task, however, presents two complications. First, it differs from a pure detection task, in which only a single stimulus (the target) is presented during each trial. Second, the target stimulus in the target-in-noise task is a drifting grating, whereas the Gaussian noise movies are distractor stimuli. This complicates the connection between our behavioral data and our electrophysiological recording data quantifying the state-dependence of V1 spiking responses to Gaussian noise movies. For these reasons, we trained a separate group of mice (N = 8 animals, n = 59 sessions) on a task in which mice were required to lick during the presentation of a Gaussian noise movie and withhold licking during a variable-length foreperiod consisting only of an isoluminant gray screen (Fig. 6a; Movie 2). In this task, therefore, the target stimuli are the same as those we used to probe the state dependence of evoked responses in V1 (Fig. 3). This task was also psychometric, with the Gaussian noise movie being presented at several contrasts during a session (Fig. 6b,c; Fig 5-1b). The state dependence of performance on this task was similar to that on the target-in-noise detection task; locomotion generally degraded perceptual sensitivity to the target Gaussian noise stimulus. For all but 1 out of 8 animals, locomotion was associated with reduced d’ compared to stillness during comparable arousal levels (Fig. 6e) and the largest d’ during stillness was significantly higher than the largest d’ during locomotion (Fig. 6f). Furthermore, for the vast majority of animals (6 out of 8), the largest d’ occurred when pupil diameter was ~30-60% constricted from the maximum (Fig. 6g). Thus, even on a visual detection task requiring only the detection of Gaussian noise movies, high arousal with locomotion significantly degraded task performance, whereas performance was comparable for a range of intermediate-arousal levels without locomotion.

One potential explanation for the apparent mismatch between the direction that high arousal/locomotion affects V1 neuronal responsiveness in passively behaving animals (increase) versus visual detection performance in task-engaged animals (decrease) is that arousal dynamics could be distinct between these two populations. Specifically, baseline pupil diameter measurements from the passively behaving versus the task-engaged population could have different latencies relative to phasic arousal events (i.e. pupil dilation/constriction and locomotion onset). It has recently been shown that the excitability of sensory cortical neurons varies during high arousal and locomotion as a function of time relative to locomotion onset (Shimaoka et al., 2018). Thus differences between passively behaving and task-engaged animals during high arousal states could be attributable to systematic differences between these populations in the temporal delay between the onset of phasic arousal events and the time at which baseline pupil diameter was measured. To test this possibility, we analyzed the distributions of latencies from pupil dilation onset, pupil constriction onset, and locomotion onset to the times at which baseline pupil diameter was measured for all recordings and animals in our passively behaving and task-engaged data set (N_passive_ = 25 animals, nmeasurements post dilation = 15,764, nmeasurements post constriction = 15,770, n_measurements_ post locomotion = 1,832; N_task-engaged_ = 20 animals n_measurements_ post dilation = 119,697, n_measurements_ post constriction = 119,739, n_measurements_ post locomotion = 8,547). We found that, while the distributions of latencies of baseline pupil diameter measurements from phasic arousal events were significantly different between the passively behaving and task-engaged population, these differences were quite small (Fig. 7-1): median latency from pupil dilation onset for passively behaving animals vs. task-engaged animals = 1.66 ± 0.02 s vs. 1.46 ± 0.005 s and K-S distance of 0.06, median latency from pupil constriction onset for passively behaving animals vs. task-engaged animals = 1.67 ± 0.02 s vs. 1.45 ± 0.005 s and K-S distance of 0.06, median latency from locomotion onset for passively behaving animals vs. task-engaged animals = 11.48 ± 0.66 s vs. 11.20 ± 0.44 s and K-S distance of 0.08. Thus, the latencies between phasic arousal events and baseline pupil diameter measurements generally differed between passively behaving and task engaged animals by no more than a few hundred milliseconds, making it unlikely that any systematic differences in these latencies between the two populations could account for the distinct effects of high arousal/locomotion in passively behaving versus task-engaged animals (i.e. enhanced sensory-evoked V1 neuronal responsiveness in the former and decreased visual detection performance in the latter).

**Figure 7-1.**
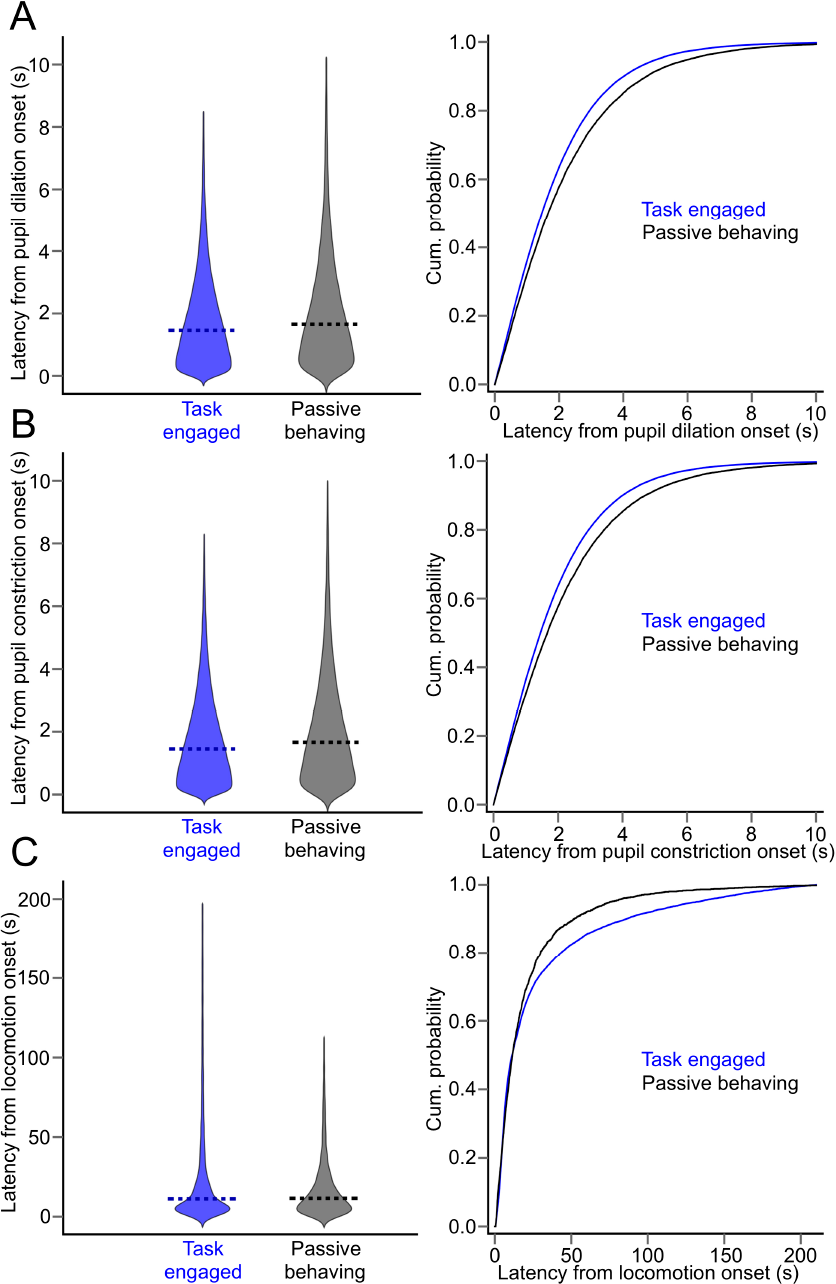
Analysis of latencies between baseline pupil diameter measurements and pupil dilation/constriction onset and locomotion onset for passively behaving and task-engaged animals. **(A)** (*Left*) Distribution charts of latencies between pupil dilation onset and pupil diameter measurements for all pupil diameter measurements made for passively behaving animals (N = 25 animals, *n* = 15,764 measurements) and task-engaged animals (N = 20 animals, *n* = 119,697 measurements). Dashed lines on charts indicate medians. (*Right*) Empirical distribution functions for the data presented at *left.* Mann-Whitney test on medians: *p ≈* 0; Kolmogorov-Smirnov test on distributions: K-S distance = 0.06, *p ≈* 0. **(B)** (*Left*) Distribution charts of latencies between pupil constriction onset and pupil diameter measurements for all pupil diameter measurements made for passively behaving animals (N = 25 animals, *n* = 15,770 measurements) and task-engaged animals (N = 20 animals, *n* = 119,739 measurements). Dashed lines on charts indicate medians. (*Right*) Empirical distribution functions for the data presented at *left.* Mann-Whitney test on medians: *p* ≈ 0; Kolmogorov-Smirnov test on distributions: K-S distance = 0.06, *p ≈* 0. **(C)** (*Left*) Distribution charts of latencies between locomotion onset and pupil diameter measurements for all pupil diameter measurements made for passively behaving animals (N = 25 animals, *n* = 1,832 measurements) and task-engaged animals (N = 20 animals, *n* = 8,547 measurements). Dashed lines on charts indicate medians. (*Right*) Empirical distribution functions for the data presented at *left.* Mann-Whitney test on medians: *p* = 0.01; Kolmogorov-Smirnov test on distributions: K-S distance = 0.08, *p* = 2 × 10^−8^.

## Discussion

Cortical state fluctuations have dramatic consequences for the processing of sensory information and the execution of sensory-guided behavior. State fluctuations in mice have traditionally been studied in the context of stillness versus locomotion, though several recent studies have begun to uncover the functional properties of state changes that occur in the absence of locomotion (Reimer et al., 2014; Vinck et al., 2015; McGinley et al., 2015a,b). Our results provide the first account of how sensory neuronal responsiveness in mouse V1 and visual detection performance vary as a continuous function of arousal. By analyzing state as a continuum, rather than by a few defined sub-states, we have shown that strong V1 sensory responses in passively behaving animals and optimal visual detection performance in task-engaged animals occur in distinct regions of the waking state spectrum (Fig. 7). Our results both complement and extend many previous results regarding the state dependence of sensory processing in the mouse visual system. We found that visually evoked neuronal spiking responses in V1 were largest and most reliable during high arousal with locomotion. This result is consistent with previous studies that have shown that locomotion is a high-gain state for V1 (Bennett et al., 2013; Dadarlat and Stryker, 2017; Fu et al., 2014; Lee et al., 2014; Mineault et al., 2016; Niell and Stryker, 2010; Pakan et al., 2016; Polack et al., 2013). Periods in which mice are still, however, encompass large and frequent state changes. Thus, without more finely analyzing the waking state spectrum, it could not be ruled out that some intermediate-arousal or non-locomotive state was in fact associated with the largest visually evoked V1 responses. We have demonstrated that this is not the case; V1 sensory responsiveness was enhanced monotonically with arousal, and largest during high arousal with locomotion (Fig. 3).

**Figure 7.**
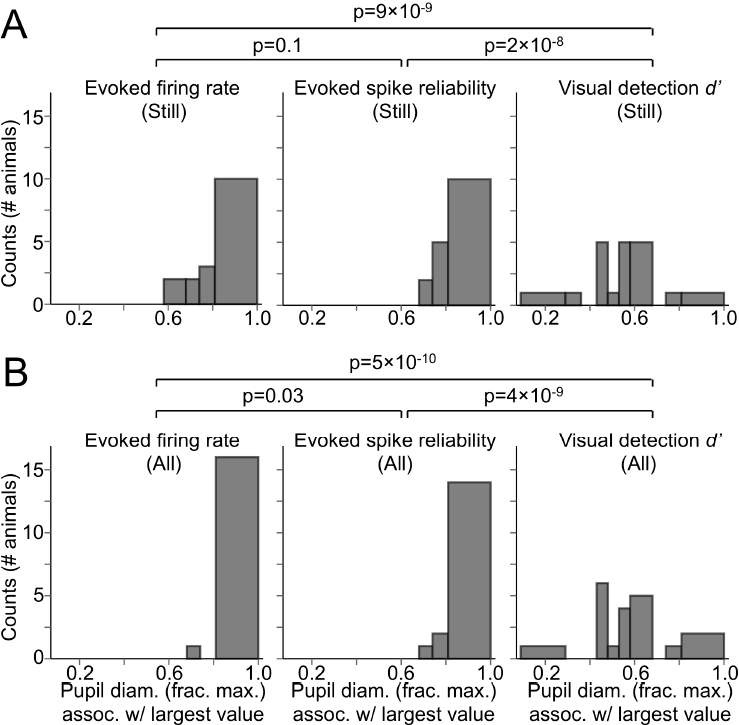
Pupil diameters associated with large and reliable visually evoked V1 responses are significantly larger than those associated with optimal visual detection performance. (A) Histograms (counts = number of animals) of pupil diameter bins during stillness associated with, left to right, largest visually evoked firing rate, spike reliability, and visual detection d’ (combined from the two visual detection tasks). Largest evoked firing rate and spike reliability recorded per animal were significantly more concentrated in bins of larger pupil diameter than the largest d’ recorded per animal (p-values from Fisher’s exact test; after Bonferroni correction, significant p-values at the 0.05 level must be lower than 0.017). For largest d’ recorded per animal, values were most strongly concentrated in pupil diameter bins ranging from 40-60% constricted from the maximal pupil diameter. **(B)** As in A, but for still and locomotion periods combined. For pupil diameter bins, bin widths were chosen such that an equal amount of data fell into each bin.

We showed that with increased arousal, V_m_ of V1 layer 5 neurons depolarized and exhibited less variability, with the most depolarized and least variable V_m_ occurring during high arousal with locomotion (Fig. 4). Several previous studies have shown that, compared with stillness as a whole, locomotion is associated with a more depolarized and less variable V_m_ in mouse V1 neurons (Bennett et al., 2013; Polack et al., 2013; Reimer et al., 2014). Our results build upon these previous data by showing that the low V_m_ variance associated with locomotion is the end point of a continual V_m_ variance reduction that occurs with increases in arousal, while enhanced depolarization primarily occurs during high arousal periods.

Many of our results highlight the notion that locomotion itself results in changes in spontaneous and evoked V1 activity beyond the changes associated with high arousal levels. Mice frequently experienced periods of high arousal without locomotion (i.e. periods in which the pupil was almost fully dilated, but without sustained locomotion bouts). We showed that locomotion, compared with still periods at a comparable arousal level, was associated with higher spontaneous V1 firing rates, larger and more reliable V1 spiking responses to visual stimuli, more depolarized and less variable V_m_, and an overall reduction in the performance of visual detection tasks. It will be important to elucidate the cellular and network basis for the additional effects of locomotion on V1 activity, even after accounting for high arousal per se. Feedback corticocortical inputs could be a prime candidate for these motor-related effects. Feedback inputs from motor cortex can exert powerful state changes in primary sensory cortical areas (Zagha et al., 2013). Thus, locomotion-dependent activation of these corticocortical afferents could somehow supplement the cortical state changes already taking place due to the enhanced neuromodulatory tone of high arousal. While mouse V1 is not known to receive direct, monosynaptic inputs from motor cortical regions, it does receive such inputs from other frontal cortical regions that might be involved in motor planning (Zhang et al., 2014; Zhang et al., 2016).

The state-dependent properties of mouse V1 activity that we have described here differ in several crucial respects from a similar analysis of mouse A1 (McGinley et al., 2015a). McGinley et al. (2015a) showed that sound-evoked A1 spiking responses exhibited maximum magnitude and reliability during intermediate arousal without locomotion. This intermediate-arousal state for augmented sensory responsiveness was associated with the most hyperpolarized and least variable V_m_ in A1 neurons. In contrast, in mouse V1, we have found that visually evoked response magnitude and reliability exhibit a continual enhancement with increased arousal, with maximal enhancement during high arousal with locomotion. This progressive enhancement with arousal was associated with a concomitant reduction of V_m_ variance in V1 neurons and depolarization during arousal, especially during locomotion. Thus, while reduced V_m_ variability is a signature for enhanced sensory responsiveness in both V1 and A1, depolarization rather than hyperpolarization is associated with large sensory-evoked responses in V1.

The mechanistic basis for the differences between V1 and A1 in the state-dependence of their sensory-evoked responses is unclear. The circuit mechanisms underlying the modulation of sensory-evoked responses by locomotion have been previously investigated in V1 and A1. In V1, a disinhibitory circuit involving inhibitory interneurons expressing vasoactive intestinal peptide (VIP) is indispensable for the enhancement of visually evoked responses by locomotion (Batista-Brito et al., 2017; Fu et al., 2014). Much of the locomotion-dependent activation of VIP cells is attributable to enhanced cholinergic tone (Fu et al., 2014), likely arising from the basal forebrain (Lee et al., 2014). In A1, on the other hand, locomotion is thought to involve an overall inhibitory circuit, in which feedback projections from secondary motor cortex (M2) predominately innervate parvalbumin (PV)-positive interneurons in A1 (Schneider et al., 2014). Importantly, however, increases in arousal from an intermediate-arousal optimum result in strongly suppressed sound-evoked responses in A1, even without locomotion (McGinley et al., 2015a). Thus, there are likely circuit mechanisms unrelated to feedback motor signals resulting in suppressed sensory responses in A1 during high-arousal states without locomotion. Generally speaking, it is worth noting that the magnitude of sensory-evoked cortical neuronal responses depend on the interplay of a number of different factors, including baseline cortical neuronal membrane potential and input resistance, the level of activation and the number of activated thalamocortical afferents, the short-term synaptic dynamics of these afferents, and the relative contribution of excitation and inhibition to the evoked cortical response. These factors appear to differ among sensory cortical areas even during the Up and Down states of slow-wave sleep and anesthesia (Haider et al., 2007; Hasenstaub et al., 2007; Neske, 2016). It is conceivable that these factors could also be distinctly modulated among different cortical areas during wakefulness.

A seemingly counterintuitive result of our study is that, despite the association between high arousal and locomotion with strong and reliable visually evoked spiking responses in V1, performance of mice on two different visual detection tasks was suboptimal at this state (Figs. 5-7). Mice generally performed our visual detection tasks best during a range of intermediate arousal levels without locomotion. The apparent mismatch between the effect of high arousal and locomotion on evoked V1 neuronal responses and visual detection performance could have depended on the spatial frequencies of the visual stimuli we used. The spatial frequencies of the visual stimuli we used in both our electrophysiological and behavioral experiments were relatively low (see Materials and Methods, “Visual stimulus presentation”). Since high arousal and locomotion afford a preferential gain enhancement to V1 neurons tuned to high spatial frequencies (Mineault et al., 2016), it is possible that visual detection task performance would be improved during high arousal and locomotion when high-acuity vision is required.

There are many possible reasons why large and reliable neuronal responses to a stimulus in a sensory cortical area might not be translated to optimal performance, even on a task that requires perception of that stimulus. First, many perceptual decisions involve the activation of many cortical areas interposed between primary sensory cortex and the motor cortical areas that generate the appropriate behavioral output. While a certain waking state might be optimal at the level of primary sensory cortex, it might not be optimal for the task-dependent neuronal dynamics of a downstream cortical area. Indeed, a long-standing hypothesis regarding the neural circuit basis for the Yerkes-Dodson principle is that during intermediate arousal, the firing properties of ascending neuromodulatory systems (in particular, noradrenergic projections) are best suited to resolving task-related neuronal dynamics in frontal cortical regions (Aston-Jones and Cohen, 2005). It will be important for future work to characterize the state-dependent processing of task-relevant sensory stimuli across the cortical processing hierarchy. Secondly, it is possible that decisions in perceptual decision making tasks, perhaps even detection tasks, are not based on a simple pooling of the sensory-evoked spiking responses of primary sensory cortical neurons. It could be, for instance, that the responses of only a certain number or class of neurons is relevant to the task. In this case, an increase in responsiveness in the entire cortical population might be detrimental to task performance, whereas a lower-gain network state might actually be beneficial (Otazu et al., 2009; Gutnisky et al., 2017). Furthermore, the perceptual decision might be based on a feature other than the summed activity of sensory cortical neurons (Montijn et al., 2015). It will be imperative for future studies to determine which features of sensory cortical responses are task-relevant (Panzeri et al., 2017), and how these features are modulated by waking state.

## Acknowledgements

We thank David Salkoff and Trevor Stavropoulos for assistance with Arduino hardware and Psychtoolbox software; Matthew McGinley and Peter O′Brien for assistance with LabVIEW programming and National Instruments hardware; Anthony Desimone for machining work. We also thank Jantine Broek, Gregg Castellucci, Matthew McGinley, David Salkoff, Paul Steffan, and Edward Zagha for helpful discussions. Support from NIH 7R35NS097287-02 (D.A.M.), NIH 1F32NS100279-01 (G.T.N.), and Yale School of Medicine Brown-Coxe Postdoctoral Fellowship (G.T.N.).

## Materials and Methods

### Animal details

For this study, male C57BL/6J mice (Jackson Laboratory) aged 7-12 weeks were used. All mice were housed on a 12-hr light/dark cycle, with ad lib access to food and water, except for water-restricted animals (see below). The light-dark cycle for all mice was reversed, such that lights in the housing facility were off from 10:00 AM to 10:00 PM and on for the remaining 12 hrs. All experiments were performed during the early-to-mid part of the dark cycle (10:00 AM - 4:00 PM). Prior to headpost surgeries (see below), mice were group-housed with 2-4 animals per cage. After head-post surgeries, mice were singly housed for ~1 week (for electrophysiological recordings) to ~1 month (for behavioral studies). All procedures involving mice were approved by the Yale Institutional Animal Care and Use Committee.

### Surgical procedures

To achieve head-fixation on the experimental apparatus, mice were first surgically fitted with light-weight aluminum headposts. Mice were anesthetized on a stereotaxic frame with 0.8–1.5% isoflurane in O_2_. After most of the scalp was resected and underlying fascia removed, the aluminum headpost was glued to the skull using dental cement (RelyX Unicem, 3M). For mice used in electrophysiological recordings, a point on the skull overlying V1 was marked prior to gluing the headpost (right V1: +2.5 (x), −3.5 (y) from bregma). After gluing the headpost, a small recording chamber was built from dental cement posterior to the base of the headpost. The recording chamber was then filled with a silicone elastomer (Kwik-Cast, WPI) to protect the exposed skull. After surgeries, mice were injected intraperitoneally with meloxicam (0.3 mg/kg) and Baytril (2.5 mg/kg) and allowed to recover on a heating pad.

To make craniotomies for electrophysiological recordings, headposted mice (by this point, habituated to head-fixation on the experimental apparatus over several days) were again anesthetized on the stereotaxic frame, and the skull was thinned with a dental drill in a ~1 mm region around the point previously marked for V1. Drilling was performed in 2-s increments followed by irrigation with cold artificial cerebrospinal fluid (ACSF) of the following composition (in mM): 135 NaCl, 5 KCl, 1 MgCl_2_, 1.8 CaCl_2_, 5 HEPES, 25 dextrose, pH 7.3 with NaOH. After the skull was sufficiently thinned to clearly visualize surface vasculature, a craniotomy of ~0.3 mm diameter was made with a microprobe (Fine Science Tools). Within this craniotomy, a duratomy as small as possible (<0.3 mm) was made with a tungsten microelectrode (FHC). A successful duratomy was evidenced by the flow of a small amount of cerebrospinal fluid. The craniotomy and duratomy were made especially small in order to prevent brain pulsations during electrophysiological recordings in awake mice. After the duratomy was made, the area was wetted with warm ACSF and silicone elastomer was applied to fill the recording chamber.

### Electrophysiological recordings

After allowing at least 1.5 hrs of recovery from craniotomy surgery, the mouse was secured on the experimental apparatus, the silicon elastomer was removed from the recording chamber, and an Ag/AgCl reference electrode was fixed in the chamber, resting on the skull. For extracellular recordings of MUA and LFP, either tungsten microelectrodes (500 kΩ impedance, FHC) or 16-channel silicon probes (item #: A1×16-3mm-50-413-A16, 50 μm spacing between recording sites, Neuronexus) were used. After drying the ACSF from the chamber, the electrode was positioned with a micromanipulator (MP-285, Sutter Instruments) to be approximately flush with the duratomy and was zeroed at this point. The chamber was then filled with ACSF and the electrode was slowly (1 μm/s) advanced into the brain. For tungsten microelectrodes, advancement was stopped at 520-550 μm, targeting V1 layer 5, while for 16-channel silicon probes, advancement was stopped at ~850 μm such that all recording contacts were in the cortex. All electrodes were left in place for at least 15 min before recordings.

For tungsten microelectrode recordings, microelectrodes were coupled to a headstage (Model 3000, AM Systems), which was further coupled to a single-channel amplifier (Model 3000, AM Systems). Extracellular signals were amplified (1000x) and filtered (0.1 Hz - 20 kHz) by the single-channel amplifier and digitized at 25 kHz with the Power1401 data acquisition system (CED) and Spike2 data acquisition software (CED). LFP and MUA signals were acquired by filtering the raw extracellular signal from 0.1 - 300 Hz and 300 Hz - 20 kHz, respectively.

For 16-channel silicon probe recordings, probes were coupled to two 8-channel 10x-gain headstages (MPA8I, Multichannel Systems), which were further coupled to a signal collector (SC8×8BC, Multichannel Systems) and signal divider (SD16, Multichannel Systems). Output from the signal divider was coupled to a 16-channel amplifier (Model 3500, AM Systems) and amplified, filtered, and digitized as with single-channel extracellular signals. All extracellular recordings (with tungsten microelectrodes and silicon probes) were performed acutely (i.e. the recording electrode was removed from the brain at the end of every session). No mouse was recorded from more than twice, and multiple recording sessions from the same mouse were always performed within a 24-hour period.

For whole-cell patch clamp recordings, borosilicate glass pipettes were pulled to final tip resistances between 4 and 8 MΩ and filled with internal solution of the following composition (in mM): 130 K gluconate, 4 KCl, 2 NaCl, 10 HEPES, 0.2 EGTA, 4 ATP-Mg, 0.3 GTP-Na, and 14 phosphocreatine-2K. Internal solutions had a final osmolality of 290-295 mosmol/kgH_2_O and pH of 7.22-7.25 with KOH. Micropipettes were secured in holders that were coupled to a CV-7B headstage (Molecular Devices), which was coupled to a MultiClamp 700B patch-clamp amplifier (Molecular Devices). Signals were filtered (DC-10 kHz) and digitized as with extracellular recordings. Whole-cell recordings were achieved using standard blind patching techniques. ~8 psi of positive pressure was applied to the micropipette before being lowered into the recording chamber bath solution. Small voltage pulses were given to the micropipette in the voltage clamp configuration. The micropipette was then lowered through the duratomy at a rate of ~15 μm/s before reaching the depth of interest (~450 μm relative to pia). At this point, pressure on the micropipette was reduced to 0.6-0.8 psi, micropipette capacitance was compensated, and the micropipette was advanced in 1 μm/s increments while monitoring the current responses to voltage pulses on an oscilloscope. Possible encounters of the micropipette with a neuron were accompanied by an abrupt increase in micropipette resistance (i.e. smaller current responses to voltage pulses). At the first sign of these resistance increases, the micropipette was advanced ~1 μm further and the positive pressure was quickly released. Cell-attached seals of >1 GΩ occurred spontaneously or through the application of a small amount of suction. Pulses of suction were then applied to rupture the neuronal membrane and achieve the whole-cell configuration, after which the series resistance was compensated (series resistances 40-60 MΩ) and the recording configuration was switched to current clamp. As reported previously (McGinley et al., 2015a), initial resting membrane potentials were markedly hyperpolarized and exhibited reduced synaptic activity, possibly due to extrusion of the high-potassium internal solution while searching for neurons. During this initial period, we recorded the intrinsic properties of the cell by stimulating the cell with 500-ms current pulses in 10-pA increments. All cells in our data set (n = 10) were classified as regular-spiking, putative pyramidal cells as evidenced by their relatively broad spike widths (mean spike width at half-height, measured from spike threshold: 0.63 ms, S.D. 0.18 ms) and rounded afterhyperpolarizations following single spikes. Within ~10 minutes of achieving the whole-cell configuration, robust spontaneous synaptic activity returned, at which point we began electrophysiological recording in combination with pupillometry and locomotion monitoring. In order to minimize possible damage to cortex, no more than 10 attempts were made at achieving a whole-cell recording per animal, and no more than 2 cells were recorded per animal.

### Visual stimulus presentation

Visual stimulus frames were presented on a 15 cm × 9 cm, gamma-corrected LCD monitor (model 665GL-70NP/HO/Y, Lilliput) (mean luminance: 20 cd/m^2^). The LCD monitor was mounted parallel with the left side of the mouse’s face, 11 cm from the left eye. The LCD monitor thereby subtended ~68° azimuth/44° elevation of the mouse’s left visual hemifield, without subtending any visual angle of the right visual hemifield or the binocular zone. A photodiode (Thorlabs, model SM1PD1B) was mounted to the upper-left corner of the LCD monitor and output to the Power1401 data acquisition board to allow precise measurements of visual stimulus onsets. Presentation of individual stimulus frames was synchronized with the 60 Hz refresh rate of the monitor using MATLAB-based Psychtoolbox software (Brainard, 1997). Visual stimuli in this study consisted of either full-contrast Gaussian noise movies, contrast-modulated Gaussian noise movies, full-contrast drifting square-wave gratings, or full-contrast drifting square-wave gratings alpha blended with full-contrast Gaussian noise movies. To construct Gaussian noise movies, individual movie frames were generated by taking the inverse Fourier transform of a 2D Fourier spectrum with a 1/(f + f_c_) decay in power, where f_c_ = 0.05 cycles per degree (cpd), and a low-pass cutoff at 0.12 cpd (Niell and Stryker, 2008). To achieve a temporal frequency spectrum with a uniform distribution from 1-4 Hz, individual Gaussian noise movie frames were alpha blended over durations randomly chosen from [1-0.25 s] such that the duration of the full Gaussian noise movie totaled 2 s (for electrophysiological recordings) or 1.5 s (for visual detection tasks). Drifting square-wave gratings used in the target-in-noise detection task (Fig. 5) had a spatial frequency of 0.05 cpd, temporal frequency of 2 Hz, and duration of 1.5 s. To modulate detection difficulty during the task, individual frames of the drifting square-wave gratings were alpha blended at varying ratios with individual frames of Gaussian noise movies. During electrophysiological recordings, 2-s long full-contrast Gaussian white noise movies were presented every ~5 s, with each inter-stimulus interval randomly chosen from an exponential distribution with a mean of 6 s and cutoffs at 5 s and 10 s.

### Habituation to head fixation and behavioral training

After allowing at least 1 day of recovery from headposting surgeries, mice were habituated by graded exposure to the experimental set-up and head fixation on a spring-mounted styrofoam wheel (20 cm diameter, 13 cm width). We first allowed mice to freely explore the wheel while they were on a small raised platform. We followed this by handling the mouse’s headpost with forceps and lifting the mouse toward the wheel. After several ~1-minute sessions of these movements, we finally secured the mouse by its headpost on the wheel for incrementally longer sessions, the first session beginning at 1 minute and later sessions lasting as long as 30 minutes. To encourage the mouse to initiate locomotion bouts during the initial habituation sessions on the wheel, we gently lifted the mouse’s tail to induce locomotion with an erect posture. Within ~2 days of habituation sessions, mice exhibited smoothly initiated voluntary walk bouts with normal posture, separated by periods of stillness.

Prior to training on visual detection tasks, mice were habituated to the experimental set-up as described above. Mice were then placed on a water restriction regimen (1 mL/day). Mice were weighed daily to ensure that their weight did not decrease below 85% of their pre-restriction baseline weight. Mice were first introduced to the lick spout on the experimental set-up as a source of water reward (i.e. free dispensation of ~10 droplets of ~4 μL). Dispensation of water was controlled by a solenoid valve under TTL control. After mice clearly recognized the lick spout as a source of water reward, training on visual detection tasks began as follows (Mice were trained 7 days/week with 2 sessions per day for the entirety of the training period and behavioral experiments, lasting ~1 month).

#### Target-in-noise detection task

On the first training session, the presentation of full-contrast drifting square-wave gratings was paired with the dispensation of one droplet of water (classical conditioning). After ~10 of these classical conditioning trials during the first session, mice proceeded to the target-in-noise detection task (Fig. 5a). At the beginning of each trial on this task, there was a variable-duration foreperiod consisting of sequences of 1.5-s long Gaussian noise movies (each movie presentation randomly chosen from 4 possible movies) (15% contrast on the simplest version of the task), with each movie separated by 0.5 s of isoluminant gray screen. The number of Gaussian noise movies presented during the foreperiod varied randomly from trial to trial: 1 to 6 movies according to an exponential distribution with a mean of 2. Licking during the Gaussian noise movies constituted a false alarm and withholding licking constituted a correct rejection (Licks were detected by interruptions of an IR beam positioned in front of the lick spout). Immediately following a false alarm, a “time-out” period began, in which the mouse had to wait 12 s before the start of another trial. During the time-out period, the screen was gray. If the mouse withheld licking during all Gaussian noise movies during the foreperiod, the 1.5-s long target period began, in which a drifting square-wave grating alpha blended with a Gaussian noise movie was presented. After an initial 100-ms period following the onset of the target stimulus, licking during the stimulus constituted a hit, and was rewarded immediately with a water droplet. Hits were followed by a 2-s long inter-trial interval (ITI) (gray screen), while misses (i.e. failure to lick during the target period) were followed by a 4-s long ITI (gray screen). Monitoring of licks and dispensation of reward were handled through digital inputs and outputs of an Arduino Uno R3 controller board (Sparkfun) and via serial communication with the behavior computer through an Arduino RS232 Shield (Sparkfun). Individual behavior sessions consisted of 400 trials, usually requiring 1.5 - 2 hrs for mice to complete. After mice maintained a d’ above 1 for at least 2 sessions for the simplest version of the task (15% contrast Gaussian noise movies, grating-noise movie alpha blending ratio of 1, i. e., full-contrast grating), the task was made incrementally more difficult for each mouse by using 50% contrast and then full-contrast Gaussian noise movies. Each mouse proceeded to the next difficulty level after maintaining a d’ above 1 for the current difficulty level for at least 2 sessions. After maintaining a d’ above 1 for the task with full-contrast Gaussian movies, mice began a psychometric version of the task, in which the drifting grating target stimuli were randomly alpha blended with one of the Gaussian noise movies with 5 different blend ratios. The first psychometric version of the task consisted of target stimuli alpha blended with Gaussian noise movies with ratios of 1, 0.84, 0.68, 0.52, and 0.36. After mice maintained a d’ above 1 for the ratio of 1 (i.e. full-contrast grating) and a d’ above 0 for the ratio of 0.68 for 2 sessions, mice proceeded to the next psychometric version of the task, with ratios of 1, 0.7, 0.5, 0.2, and 0.05. After performing with a d’ above 0 for a ratio of 0.5 on this task for 2 sessions, mice reached the final psychometric version of the task, consisting of blend ratios 1, 0.32, 0.08, 0.02, and 0.005. Most mice reached the final psychometric version of the task within ~2 weeks of the first training session (~28 sessions). Mice were trained on the final psychometric version of the task for 2 days (4 sessions) before we recorded pupillometry and locomotion data during task performance. During sessions with pupillometry and locomotion data collection, target stimuli with blend ratios of 1, 0.32, and 0.08 were presented.

#### Noise detection task

The training procedure and trial structure for the noise detection task were essentially identical with those of the target-in-noise detection task, with a few key differences (Fig 6a). For the noise detection task, the target stimulus was a single Gaussian noise movie, presented at different contrasts during the session. Trial onset was signaled by a 200-ms audible tone, after which a variable-length foreperiod began (duration of 2-12 s, exponentially distributed with a mean of 4 s). This foreperiod consisted only of a gray screen, and a correct rejection was recorded every 1.5 s that the mouse withheld licking, while licking during this foreperiod constituted a false alarm. The simplest version of the task consisted of only the full-contrast Gaussian noise movie as the target stimulus. As mice reached performance criteria as with the target-in-noise detection task, psychometric versions of the task were made progressively more difficult, starting with Gaussian noise movie contrasts of 100%, 84%, 68%, 52%, and 36%, progressing to 100%, 70%, 50%, 20%, and 5%, and ending with the final psychometric version of the task with 100%, 40%, 10%, 5%, and 1%. During sessions with pupillometry and locomotion data collection, Gaussian noise movies of 100%, 10%, and 5% were presented.

### Locomotion monitoring and pupil video acquisition

Locomotion speed was recorded by mounting an optical mouse (Logitech G600) next to the wheel. USB output from the optical mouse was converted to an analog signal proportional to wheel speed using custom-written LabVIEW (National Instruments) software. The analog signal was output by a small DAQ device (USB-6008, National Instruments) to the main Power1401 data acquisition system.

To acquire video of the mouse’s right eye, a USB camera (Basler ace acA1300-30um) equipped with a C-mount lens (Basler Lens C125-1620-5M F2.0 f16mm) and 780 nm long-pass filter (Thorlabs FGL780M) was positioned ~11 cm from the right side of the mouse’s face, which was illuminated by 2 IR LEDs. The visible light from the LCD monitor was sufficient to allow an appropriate dynamic range for state-dependent pupillary fluctuations. Video capture from the camera (~30 Hz frame rate) was controlled by custom-written LabVIEW software. Video frames were 1280×960 pixels. To synchronize video frames with other data acquired during the experiment, an aperiodic train of digital pulses from an Arduino was simultaneously sent to the main Power1401 data acquisition system and an additional small DAQ device (USB-6008). The LabVIEW software synchronized video frames with the digital pulse train read in by the small DAQ device by updating the intensity of a collection of pixels on the frame according whether the current value of the pulse train was 0 V or TTL+.

### Atropine experiments

In one cohort of mice (N = 17 animals), we applied atropine sulfate ophthalmic solution (1%) (Akorn) to the left eye prior to electrophysiological recordings to keep the pupil of the left eye maximally dilated for the entire recording (Fig. 3-2). After allowing at least 1.5 hrs of recovery from the craniotomy surgery, the mouse was secured on a separate set-up from the main electrophysiology rig. On this set-up, two pupillometry cameras were mounted to capture simultaneous video of the left and right eye. We first took a baseline video, which showed coherent pupillary fluctuations in the left and right eye. While the mouse was still secured on this set-up, we applied 30 μL of the atropine solution to the left eye, and allowed the mouse to recover in its cage for an additional hour. We then placed the mouse back on the double-pupillometry set-up and took a video of the left and right eye to verify that the left pupil was completely dilated while the right pupil fluctuated with state changes. We then secured the mouse on the main electrophysiology rig for recording. After an electrophysiological recording experiment, we placed the mouse on the double-pupillometry set-up a final time to ensure that the left pupil was still completely dilated, and was thus completely dilated during the electrophysiological recording experiment.

### Experimental Design and Statistical Analysis

In the both the text and figures, “N” refers to the number of animals and “n” refers to the number of recordings (extracellular), cells (whole-cell), or sessions (behavior experiments). Unless otherwise noted, data are summarized in both the text and figures as the mean ± 68% bootstrap confidence intervals. For electrophysiology data, the mean represents the mean over all recordings or cells (no more than 2 recordings or cells per animal). For behavioral data, the mean represents the mean across animals. With the exception of data in Figs. 7, 7-1, and 3-2, we used non-parametric paired statistical tests: the rank-sum test for 2 paired variables. Paired comparisons were made within a recording for electrophysiological data and within an animal for behavioral data. For Figs. 7 and 3-2, we used Fisher’s exact test on contingency tables for pupil-diameter-binned animal counts. For Fig. 7-1, we used the Mann-Whitney test and the two-sample Kolmogorov-Smirnov test for distributions of passively behaving and task-engaged animals. P-values <0.05 were considered statistically significant. Bonferroni correction was made for P-values resulting from multiple comparisons, with a corrected cutoff of <0.05/ncomparisons.

### Quantification of pupil size

Pupil diameter was estimated from individual video frames. Frames were first cropped to regions-of-interest around the eye. Pixels in the resulting frames were then binarized such that the majority of pixels within the pupil were black and the majority of pixels outside the pupil were white. Canny edge detection with a pixel radius of 10-15 was then used on the binarized images to detect the edges of the pupil. Pupil diameter was then estimated as the authalic diameter of the shape formed by the detected edges (i.e. the diameter of a circle with the same surface area as the shape). To facilitate pooling of data across recordings and mice, raw pupil diameters in units of pixels were normalized to the value of the largest pupil diameter imaged during the session, which always occurred when the mouse was engaged in sustained locomotion bouts. Traces of normalized pupil diameter were low-pass filtered at 3 Hz to minimize any movement or eye blink artifacts.

For analysis of latencies between pupil dilation and constriction onset and baseline pupil diameter measurements (Fig. 7-1), traces of normalized pupil diameter were low-pass filtered at 1 Hz and differentiated. Points at which the derivative changed from negative to positive or positive to negative were considered onsets of pupil dilation or constriction, respectively.

### State-dependent electrophysiological metrics

We studied the dependence of spontaneous and evoked activity in V1 as a function of state by analyzing several electrophysiological variables as a function of baseline pupil diameter We also sorted data according to whether the mouse was walking or still. In order to be classified as a walking period, the average wheel speed during that period must have exceeded 5 cm/s. Furthermore, in order to be considered a still period, no walking periods must have occurred for at least 2 s prior to or following that period.

Most of our recordings were from V1 layer 5, though we also present a smaller amount of layer 2/3 data. When using 16-channel silicon probes (see above), we estimated the boundaries of cortical layers using current-source density (CSD) analysis (method described in Higley, 2012) of the LFP responses to 50-ms full-screen flashes. Layer 4 was considered to be at the depth of the mid-probe contact for which a large, short-latency CSD sink was recorded (Fig. 3-1a). Contacts 50-100 μm above the layer 4 contact were considered to be in layer 2/3.

For each extracellular or whole-cell recording, we first computed the cross-correlogram between pupil diameter and multi-unit firing rate or V_m_ (Multi-unit spikes were defined as events that exceeded 4× the standard deviation of the noise floor in extracellular signals filtered from 300 Hz - 20 kHz). We performed this initial step to determine the temporal lag between changes in cortical activity and changes in pupil diameter. For MUA and whole-cell recordings, pupil diameter changed 1.5 ± 0.1 s and 1.3 ± 0.2 s after a corresponding change in firing rate or V_m_, respectively. This lag likely exists due to the relative rapidity with which changes in neuromodulatory tone affect cortical activity versus smooth muscle contraction in the eye.

To analyze spontaneous multi-unit firing, we examined 500-ms regions occurring 2 s before a visual stimulus presentation, binned the multi-unit spikes in 20-ms windows, and divided the PSTH by 0.5 s to calculate firing rate in Hz. The pupil diameter associated with a spontaneous firing rate in a given 500-ms region was considered as the pupil diameter averaged over 500 ms, but shifted in time according to the pupil diameter-cortical activity lag determined for that recording. We note, however, that not accounting for this lag in our analysis did not affect the fundamental relationships between baseline pupil diameter and V1 cortical activity that we observed (Fig. 2-1), likely due to the slow (<1 Hz) time-course over which state and pupil diameter fluctuate. For each recording, the spontaneous firing rates associated with all the analyzed time intervals were normalized to the largest spontaneous firing rate recorded. Pupil diameters associated with each analyzed region were then sorted into bins. These bins were determined by dividing the range of pupil diameters recorded during the session into 9-11 bins, with an equal amount of data in each bin. Thus, the width of the bins at or near the extremes of the recorded pupil diameter range were often larger than those occurring at mid-range. The spontaneous firing rates in each of the 9-11 pupil diameter bins were then averaged to generate spontaneous firing vs. pupil diameter data for that recording.

To analyze MUA evoked by the presentation of Gaussian noise movies, PSTHs using 20-ms bins were constructed for the 2 s during stimulus presentation and the 500-ms period just before stimulus presentation. In a given recording session, the same Gaussian noise movie was presented repeatedly. Evoked firing rate as a fraction of baseline firing rate was calculated by dividing the firing rate during the first 1 s of the stimulus by the firing rate in the 500-ms baseline period just before stimulus presentation. The pupil diameter associated with a given evoked firing rate was calculated as with spontaneous periods, described above. Evoked firing rates were also assigned to pupil diameter bins and averaged as with spontaneous periods.

The trial-by-trial spike reliability associated with a given pupil diameter bin in a recording was determined by calculating the pairwise cross-correlation between all evoked 1-s PSTHs in that bin. To correct for increases in cross-correlation due to increases in firing rate alone, we also calculated a “chance” cross-correlation measure, which was the average pairwise crosscorrelation between each evoked 1-s PSTH and all spontaneous 1-s PSTHs associated with that pupil diameter bin.

For whole-cell recordings, spikes were first removed from V_m_ traces by filtering the traces with an 8-ms median filter. Coherence between pupil diameter and V_m_ was computed as |*P*_*pupii-Vm*_ (*f*)|^2^/(*P*_*Pupll-Pupii*_(*f*)* *P*_*Vm-Vm*_(*f*)), where *P*_*pupll-vm*_ (*f*) is the cross-spectral density between pupil diameter and V_m_, *P*_*pupll-pupii*_(*f*) is the auto-spectral density of pupil diameter, and *P*_*Vm-Vm*_(*f*) is the auto-spectral density of V_m_, all estimated using Welch’s overlapped averaged periodogram method. Spontaneous average V_m_ as a function of pupil diameter was calculated as with spontaneous multi-unit firing rates, except that each 20-ms bin of the PSTH represented the average V_m_ in that bin. V_m_ variability during a given spontaneous period was calculated as the standard deviation of the V_m_ during that period. To obtain the 2-10 Hz and 50-100 Hz Hilbert amplitude of V_m_ traces, we first bandpass filtered V_m_ traces at 2-10 Hz and 50-100 Hz and then computed the complex conjugate of the Hilbert transform on the filtered traces, representing the instantaneous amplitude envelope of these traces. Though we presented Gaussian noise movies during all whole-cell recordings, most cells exhibited only weak postsynaptic potentials in response to the visual stimuli, and for most cells the signal-to-noise ratio of the evoked postsynaptic potentials was below 1. Thus, we did not analyze evoked postsynaptic potentials as a function of pupil diameter. All absolute V_m_ values reported were corrected for a 14 mV liquid junction potential.

### State-dependent behavior metrics

Data from performance on visual detection tasks were analyzed on a per-animal basis. Hits, misses, false alarms, and correct rejections were pooled across all the sessions that an individual animal performed while pupillometry and locomotion data were collected (5-8 sessions per animal). Each hit, miss, false alarm, and correct rejection had an associated baseline locomotion speed value and baseline pupil diameter value, calculated as the average locomotion speed and pupil diameter during the 500 ms of isoluminant gray screen presentation before the behavioral event occurred. As with analysis of electrophysiological data, the pupil diameter range for each animal was divided into 9-11 bins, and hits, misses, false alarms, and correct rejections were assigned to pupil diameter bins based on their associated baseline pupil diameter values. For each pupil diameter bin for each animal, the hit rate was calculated as, hit rate = total # hits / (total # hits + total # misses), and false alarm rate was calculated as, false alarm rate = total # false alarms / (total # false alarms + total # correct rejections). Perceptual sensitivity (d′) for each pupil diameter bin was calculated as d = norminv(hit rate) - norminv(false alarm rate) and decision bias (c) was calculated as c = −(norminv(hit rate) + norminv(false alarm rate))/2, where norminv is the inverse of the cumulative normal function.

***Movie 1.*** Real-time performance of a mouse on different trial types during the target-in-noise detection task.

***Movie 2.*** Real-time performance of a mouse on different trial types during the noise detection task.

